# Learning robust gene expression embeddings for disease subtyping, biomarker identification and cross-species alignment

**DOI:** 10.1101/2025.01.03.631230

**Authors:** Xin Chen, Kathleen M Smith, Yingtao Bi

## Abstract

High dimension gene expression measurements such as microarray and RNA-seq data, are often plagued by sources of unwanted variation. This variability can lead to the obscuring of meaningful biological signal by technical noise and non-interesting biological variation, thus resulting in failure to identify the same set of targets and biomarkers in independent studies. This phenomenon contributes to the so-called reproducibility crisis and makes preclinical drug development more challenging. There is an urgent need for an improved process to identify shared biological signals in large datasets from independent cohorts. In this article, we propose an innovative method called Joint-Embedding via Canonical Correlation Analysis (JECCA), which aims to capture relevant biological variation by learning of shared lower dimension embedding across multiple datasets.

We demonstrate the efficacy of JECCA in analyzing multiple human cancer and inflammatory disease gene expression datasets, showcasing its ability to identify robust, biologically relevant signals. By leveraging these embeddings, our approach enables several important applications: (1) robust disease endotype identification by joint unsupervised analysis of multiple independent gene expression datasets (2) reproducible and biologically meaningful biomarker identification to enhance prediction of treatment outcome (3) cross-platform analysis, such as integration of RNA-seq and microarray data, which are accumulating exponentially in public databases (4) cross-species analysis to identify conserved molecular features between human and mouse. We provide comparative analyses that highlight the superior performance of JECCA over other approaches such as batch value-corrected methods based on joint analysis. As analysis of publicly available datasets becomes more prevalent, we foresee JECCA being employed in multiple disease contexts.

## Background

The identification of disease driver genes, discovery of robust molecular disease endotypes, and exploration of biological meaningful biomarkers from high-dimensional omics data is critical for understanding disease etiology and predicting clinical outcomes. However, achieving robustness and performance on independent datasets is challenging due to multiple factors, including random noise, biological disease heterogeneity, difference in sample collection and processing procedures^1,2^, subject characteristics including gender, age, and medications usages^3^ and small sample sizes in individual studies. These challenges often lead to masking of biologically meaningful signal within the data, rendering features learned from individual gene expression datasets nongeneralizable^4^. For example, conventional principal component analysis (PCA) is a commonly used dimension reduction method for analysis of gene expression data. Often, the first several PCA dimensions presented in a single study fail to detect the desired biological signal of interest when the effect size is small, as with peripheral blood mononuclear cell (PBMC) samples^5^.

While meta-analysis, by combining results of multiple studies, is useful to derive pooled estimates in a supervised setting, it is less powerful and ignores important differences between heterogeneous studies. It is also inadequate for unsupervised analysis. Combining independent but related datasets for joint analysis holds promise for increasing statistical power and generalizability. Often, such integration is hindered by the presence of batch effects and other sources of dataset specific variation. Although various methods^6–8^ have been developed to reduce this undesirable variability before integrative analysis, their effectiveness can be compromised by confounding factors hidden in the true biological signals^9^. When the degree of confounding between the batch variable and the endpoint of interest is high, all current methods carry the risk of removing relevant biological signal which reduces our ability to extract underlying patterns^10–12^. Moreover, most batch correction methods make parametric assumptions that are often invalid in biological realistic scenarios, potentially leading to unreliable results when clustering or building classifiers directly on batch corrected values.

Another challenge in integrative analysis arises when independent data come from different platforms or even different species, leading to an out-of-distribution problem. Cross-species alignment analysis is essential in drug development research to validate results from in vivo animal studies to transition findings to human. Recently a predictive tool called TransComp-R was developed to project human pre-treatment transcriptomic data onto the PCA variations built from mouse proteomic data. The aim was to identify predictive features of patient response to therapy by assuming that discernable variations in the mouse data are useful for building predictive models for human data^13^. However, TransComp-R was designed to integrate a single human dataset and a single mouse dataset, not to identify shared signal across multiple datasets. The direct translation of the principal components between mouse and human in TransComp-R might also be negatively impacted by the out-of-distribution issue. In a recent review paper, Foltz et al showed that some batch correction methods are useful to merge gene expression datasets across platforms for machine learning tasks when large data sets of samples are available from both platforms^14^. Methods which are useful to translate relationships across species remain limited^15^. In addition, several methods have been proposed to integrate gene expression data between cell lines and human tumor profiles, but it is not clear whether they apply to integration of more than two gene expression datasets which would increase the robustness of downstream analysis^16–18^.

In this article, we introduce an innovative approach, called JECCA (https://github.com/xchen004/JECCA), to address the challenge of extracting and aligning common biological meaningful signals across independent datasets. Inspired by recent successful efforts in integrative single cell RNA-seq data analysis to identify common cell types^19^, JECCA aims to identify shared embeddings that represent true biological variations across multiple datasets, focusing on signals independent of experiment specific technical factors. The method assumes that signals shared by multiple datasets more likely reflect true biological variations, such as representing cell type composition variations.

To demonstrate the performance of JECCA, we applied the method to several different case studies, highlighting its capabilities in different applications. First, we showed the ability of JECCA-derived embeddings to perform robust breast cancer subtyping. Second, in ulcerative colitis, JECCA generated conserved biological embeddings resulting in disease subtyping and identification of robust biomarkers to improve TNF-alpha inhibitor response prediction accuracy. Third, we successfully identify biological signals in PBMC samples for rheumatoid arthritis (RA) using JECCA-derived embeddings, despite the weaker signal present in blood derived samples. Finally, JECCA enabled cross-species analysis revealed conserved molecular features between human and mouse.

## Results

JECCA does not aim to correct the data and remove batch effects but instead identifies shared variations and their contributing gene signatures, which differs from the data-merging methods based on batch effect correction since those methods analyze the pooled data after removing eventual biases^20,21^. The flowchart of JECCA is described in Figure 1 to integrate two gene expression datasets, X and Y. The data integration strategy used in JECCA is based on Canonical Correlation Analysis (CCA), which has been used extensively to integrate multi-omics datasets which were profiled on the same set of samples^22,23^. To our knowledge, CCA has not been used to integrate multiple bulk gene expression datasets. In this article, we have twisted conventional CCA as a dual form to integrate multiple gene expression datasets which have the same set of genes measured. Those shared embeddings identified by our method have increased likelihood to be associated with relevant biological signals, which can be used for downstream analysis.

**Figure 1.**
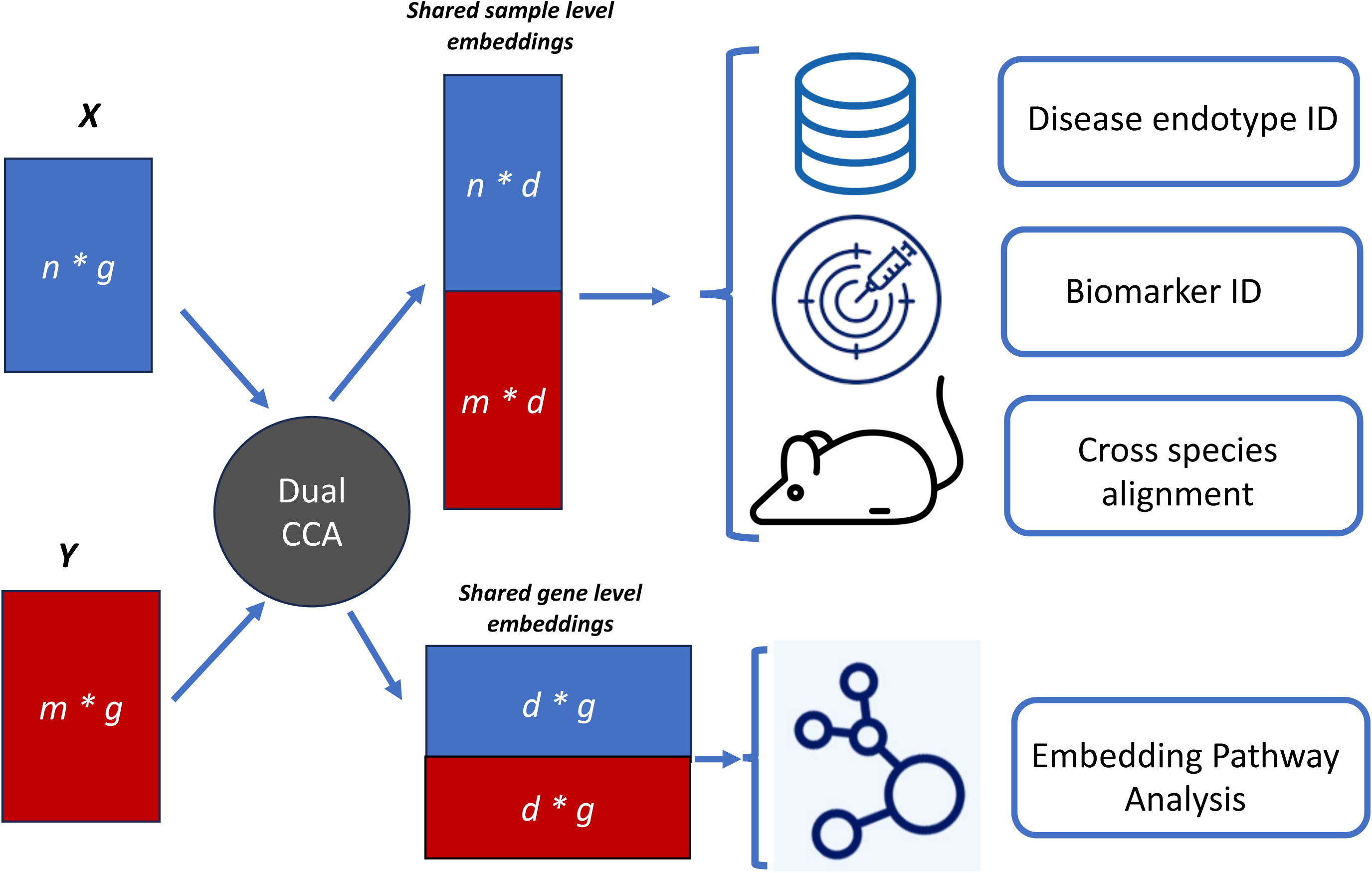
Overview of JECCA and applications. To integrate two gene expression datasets *X* and *Y*, dual CCA is applied to compute the canonical variables. Each of the original gene matrices *X* (with *n* samples and *g* genes) and *Y* (with *m* samples and *g* genes) is decomposed into two matrices, the shared sample level embeddings (with *d* embeddings), which can be used to perform downstream integrative analysis, and the shared gene level embeddings, which can be used to identify corresponding biological pathways.

### Breast Cancer Subtype Classification

For endotype identification in breast cancer, we utilized breast cancer datasets integrated in a recently published paper^24^, which analyzed 11 microarray studies comprising 1604 samples. The disease subtypes in this study were defined based on estrogen receptor (ER) status or cancer grade. We applied JECCA to the expression data to generate embeddings and performed unsupervised clustering. The UMAP plot (Figure 2A) demonstrated that the data from the 11 studies were well-mixed, indicating that the JECCA embeddings were not impacted by the individual studies. This is expected since JECCA only captures shared information across studies. Furthermore, the UMAP plots (Figure 2B-D) colored by biological variables of interest revealed that the clusters were strongly associated with ER status and cancer grade. Cluster 2 (Fig 2B-C), for example, exhibited an enrichment of samples devoid of ER positive and graded as 3. Within cluster 2, 75.9% of the samples were labeled as ER negative, while the remaining clusters had only 14.1% labeled as ER negative (p-value = 8.3e-71 in a Chi-squared test between ER status and identified cluster). Similarly, 72.7% and 46.5% of the samples in cluster 2 and cluster 1, respectively, were labeled as grade 3, compared to only 12.8% in the other clusters (p-value = 4.5e-52 in a Chi-squared test between cancer grade and estimated cluster).

**Figure 2.**
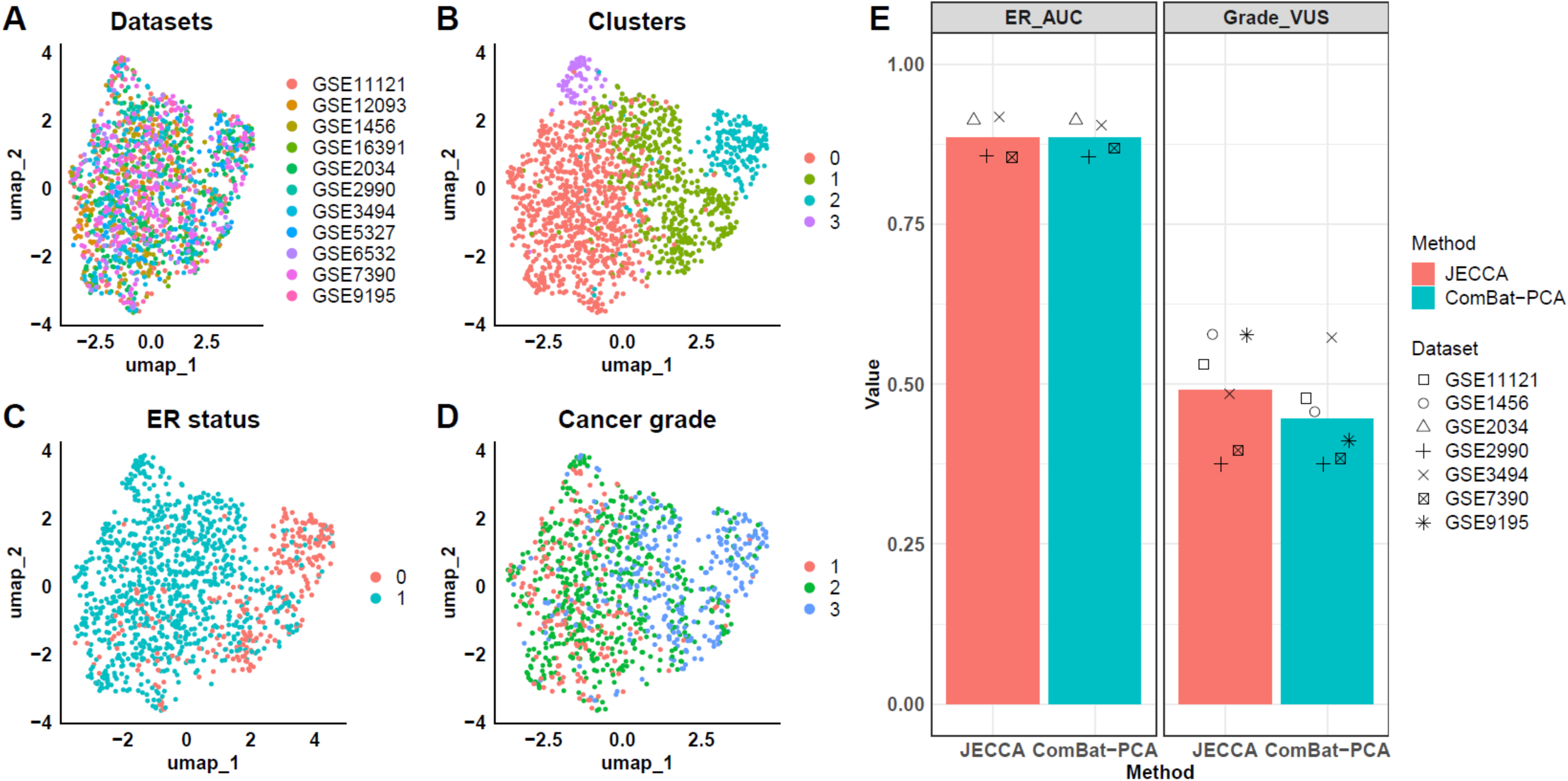
Breast cancer clustering and classification analyses. (A-D) UMPA plots of JECCA-derived embeddings. Plots are colored by (A) GEO datasets, (B) Estimated clusters, (C) ER status, and (D) Cancer grade. (E) Barplots of AUC and VUS estimated in ER status and cancer grade classification analyses. In classification analysis, the ten breast cancer datasets were used as training data and another one dataset was used as testing data.

To further assess the predictive power of the JECCA-derived embeddings, we employed the embeddings from analysis of the 10 datasets as training data and reserved one for testing. We conducted classification analysis to predict ER status and cancer grade. Random forest was employed to build a classifier on the training data and predict the ER status and cancer grade on the testing data. Due to the limited numbers of several defined subtypes in some of the datasets, we selected specific datasets for testing to ensure robust classification analysis. For ER prediction, we used GSE2034, GSE2990, GSE3494, and GSE7390 as testing data. For cancer grade prediction, we used GSE1456, GSE2990, GSE3494, GSE7390, GSE9195, and GSE11121 as testing data to make sure each testing dataset has enough sample size for each subtype. The performance of the classifier was evaluated using the area under the ROC curve (AUC) and the volume under a three-class ROC surface (VUS). VUS is a metric that expands upon the concept of AUC for three-class classification problem. Similar to the AUC metric, a higher VUS value suggests superior classification performance. To illustrate, a VUS value of 1 represents a perfect classifier, while a value of 1/6 is indicative of random chance classification.

For comparison, we applied ComBat, which is the most widely used method for batch effect correction, on the 11 datasets to remove batch effects and performed principal component analysis (PCA) on the batch-corrected data to generate competitor embeddings. Both VUS and AUC (Figure 2E) indicated that the JECCA embedding classifier outperformed the ComBat-PCA embeddings classifier in cancer grade prediction and performed as well as ComBat-PCA in ER status prediction. The clustering and classification results demonstrated that the JECCA embeddings preserved shared biological information associated with breast cancer subtypes defined by ER status and cancer grade.

### Robust Disease Subtyping and Biological Meaningful Biomarker Identification in Inflammatory Bowel Disease (IBD)

Next, we turned our attention to autoimmune disease ulcerative colitis (UC), wherein most patients with public data have been treated with anti-TNF therapies such as adalimumab (ADA) or infliximab (IFX). While clinical trials suggest that patient response as measured by clinical remission is 16.5-37% depending on the drug and the study, real life data shows that the overall response rate to these therapies is often poor due to disease (or patient) heterogeneity^25–27^. Thus, it is extremely important to identify robust biomarkers to assess anti-TNF therapeutic efficacy for optimal personalized treatments and improve our understanding of disease pathophysiology.

For this analysis we focused on four microarray studies^28–31^ of UC, encompassing a total of 102 patients treated with the anti-TNF therapy IFX. Czarnewski et al. used these same four datasets and showed that standard gene-ranking strategies do not allow clustering of UC patients into consistent molecularly district subtypes without incorporating knowledge learned from murine model data^32^. We applied JECCA to integrate these same four datasets. Our analysis of the baseline samples was like that described for the breast cancer application. Similar to the breast cancer study, a UMAP plot was generated to visualize the JECCA embeddings. Figure 3A clearly shows the samples from different datasets are fused, as expected of JECCA. Clustering analysis (Figure 3B, 3C) revealed two distinct clusters significantly associated with IFX responder/non-responder status (p-value = 0.030 in a Chi-squared test).

**Figure 3.**
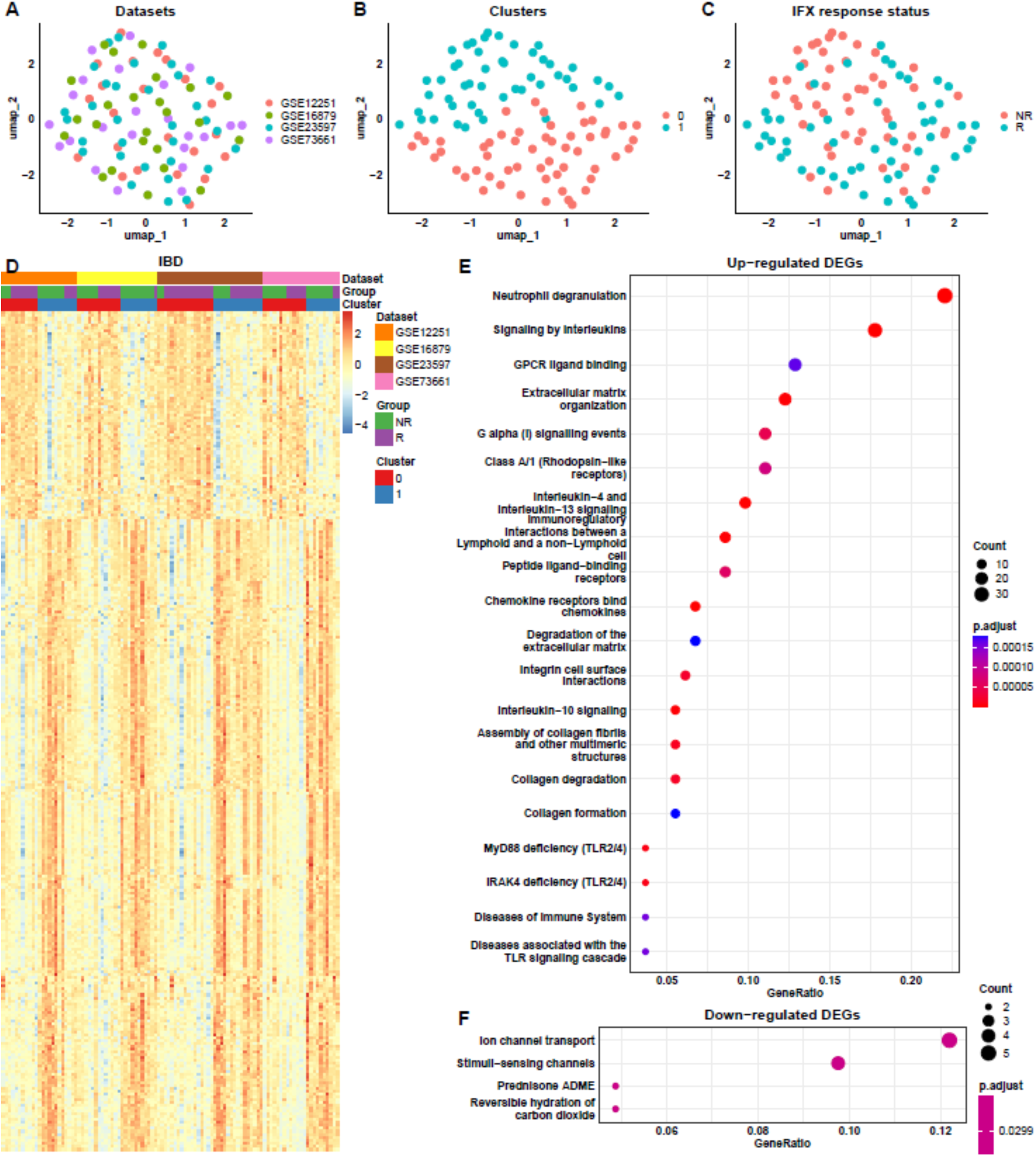
UC samples clustering and DEG analysis. (A-C) UMPA plots of JECCA-derived embeddings. Plots are colored by (A) GEO datasets, (B) Identified clusters, and (C) Response status to IFX. (D) Heatmap of DEGs identified between the two clusters (cluster “1” vs cluster “0”). Samples are annotated by the estimated clusters, response status to IFX, and GEO datasets. (E-F) Pathway analysis of (E) upregulated and (F) downregulated DEGs using the reactomPA database. Significant pathways are selected based on adjusted p-value < 0.05 and a cutoff of top 20 pathways.

Next, we conducted a differential expression analysis between the two clusters (cluster “1” vs cluster “0”), identifying 295 differentially expressed genes (DEGs) comprising 222 up-regulated and 73 down-regulated genes. The heatmap (Figure 3D) displayed distinct gene expression profiles between the two clusters and the DEGs have consistent pattern across all datasets. Subsequent pathway analysis (Figure 3E) of the up-regulated DEGs revealed many interesting and previous known associations, such as Neutrophil degranulation, Extracellular matrix organization, IRAK4 deficiency, and Interleukin-4 and Interleukin-13 signaling. IRAK4 has been considered as a therapeutic target for treating ulcerative colitis^33^ and Interleukin-4 (IL-4) and IL-13 are key immunoregulatory cytokines, whose dysregulation may contribute to a range of inflammatory disease states, including ulcerative colitis (UC), asthma^34^, and atopic dermatitis^35^.

To further explore biological information preserved in each JECCA embedding, we conducted pathway analysis on the gene expression profiles associated within each JECCA embedding. We utilized GSEA-GO analysis (Figure 4A), which revealed significant pathways related to mitochondrial dysfunction, extracellular matrix, and antimicrobial activity. Our focus was particularly on three embeddings: JECCA1, JECCA3, and JECCA10, as all correlate significantly with the IFX responder status (Supplementary Table 2). Notably, JECCA1 and JECCA10 showed reverse patterns of GSEA scores for most pathways. JECCA1 embedding was highly enriched in the extracellular matrix, specifically collagen-containing extracellular matrix, while JECCA3 embedding was highly enriched in antimicrobial activity, including secretory granule, secretory granule membrane, and ficolin-1-rich granule. GSEA-ReactomPA results (Figure 4B) confirmed the findings and further identified the IL-1 family signaling pathway in JECCA3. These pathway results suggest that JECCA1 is closely associated with stroma activity, while JECCA3 is highly associated with neutrophil activity. Our findings align with a recently published paper^36^ that identified similar pathways enriched in UC stromal cells, namely neutrophils, extracellular matrix, and antimicrobial activity by performing gene co-expression module analysis. The paper identified modules associated patient therapy response to anti-TNF and verified that their high expression represents high neutrophil infiltration and activation of fibroblasts through further scRNA-seq experiments. They further demonstrated that this novel stroma-neutrophil interaction was driven by IL-1.

**Figure 4.**
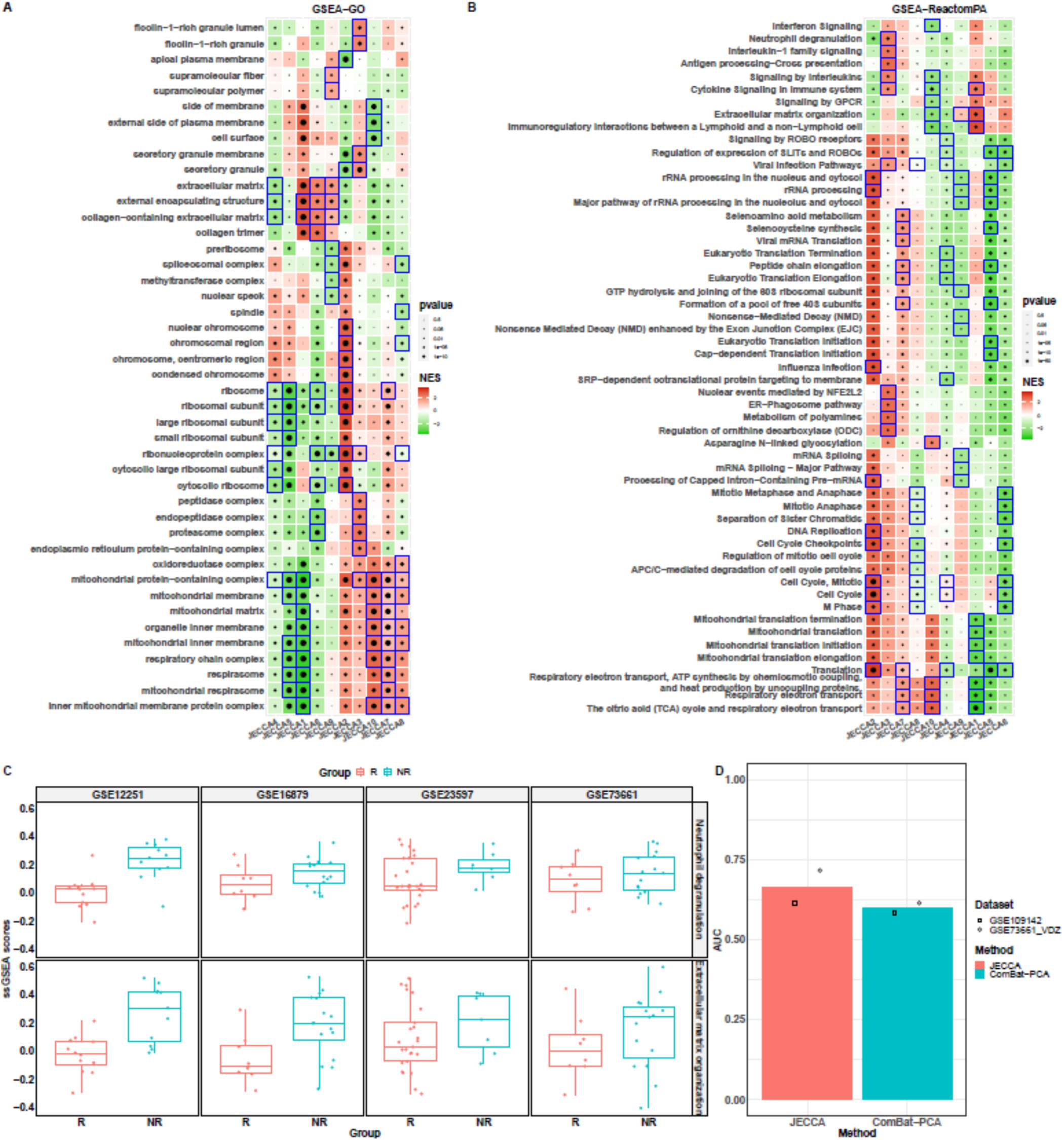
Biological interpretation of JECCA embeddings and classification analysis in ulcerative colitis study. (A-B) GSEA of 10 JECCA embeddings using (A) ‘Cellular component’ Gene Ontology database and (B) ReactomPA database. Blue boxes highlight the top 10 significant pathways in each embedding. (C) Boxplot of ssGSEA scores of two important pathways between IFX responders and non-responders across four IBD datasets. (D) Comparison of JECCA and ComBat-PCA methods by classification analysis on six datasets. In classification analysis, the four IBD human datasets were used as training data and the two additional datasets were used as testing data.

To further demonstrate the robustness of the biological signals identified by JECCA, we performed ssGSEA analysis on two interesting pathways, neutrophil degranulation, and extracellular matrix organization. The ssGSEA scores shown in Figure 4C clearly indicate that both pathways have higher expression in non-responders than responders in each dataset consistent with these recent findings. To summarize, JECCA embeddings are robust and biological interpretable. Their easy interpretation is preferable than deep learning-based approaches^21^, which usually require high end infrastructure to train and is limited by model interpretability, which is especially important for biological data analysis.

The strength of the JECCA method lies in its ability to extract shared information across multiple datasets while effectively removing uninformative factors that are not shared, such as technical artifacts and biological confounders. To demonstrate that the shared embeddings can perform well across platforms and with different drug treatments, we extended its application to a case study using the previously included four UC datasets as training data and two additional testing datasets. The two testing datasets represented distinct cohorts, one RNA-seq data from pediatric UC patients treated with IFX^37^, and one microarray RNA-seq data consisting of adult UC patients treated with vedolizumab (VDZ)^29^. We applied both JECCA and ComBat-PCA to the six datasets and conducted classification analysis to predict responder status. Figure 4D demonstrated that the JECCA method achieved better AUC compared to the ComBat-PCA.

To summarize, in this application, by jointly analyzing multiple UC gene expression datasets using JECCA, we demonstrated that JECCA-derived embeddings can be utilized to identify robust distinct molecular UC subtypes with different cellular signatures and significantly different response rates to anti-TNF therapy. Additionally, we designed an approach to identify gene biomarkers by combining JECCA gene embeddings. We showed that JECCCA embeddings identify interpretable biology and gene set enrichment analysis of the embeddings revealed robust biomarkers across the datasets.

### Prediction of Clinical Response to anti-TNF treatment in Rheumatoid Arthritis Blood Samples

Blood-based gene expression biomarkers to predict drug response are the gold standard for precision medicine. However, biomarker identification from either PBMC or whole blood samples is challenging since the biological signal is weak and impacted by differences in the proportion of immune cell subtypes. In recent studies^38,39^ gene biomarkers in blood were proposed to predict response or non-response to anti-TNF therapies in rheumatoid arthritis (RA) patients. These proposed biomarkers either lacked validation or exhibited poor prediction power in independent studies, suggesting limited robustness across different RA cohorts^40,41^. Considering this, we aimed to further demonstrate the robustness of the JECCA by utilizing it to predict responders to anti-TNF therapy in RA patients using blood samples. We incorporated data from six RA studies^39,42–45^ where patients were treated with infliximab, adalimumab (ADA), or rituximab (RTX). Moreover, these datasets were generated using various technical platforms, including RNA sequencing (RNA-seq) and three different types of microarrays. In addition to including ComBat-PCA for comparison, we also employed the quantile normalization method, which performed best for combining microarray and RNA-seq data for supervised machine learning in a recent review study^46^. Similar to ComBat-PCA, we performed quantile normalization on the expression data and subsequently applied principal component analysis (PCA) to generate low dimensional embeddings. The results of the classification analysis revealed that the JECCA outperformed both the ComBat-PCA and QN-PCA methods (Figure 5A).

**Figure 5.**
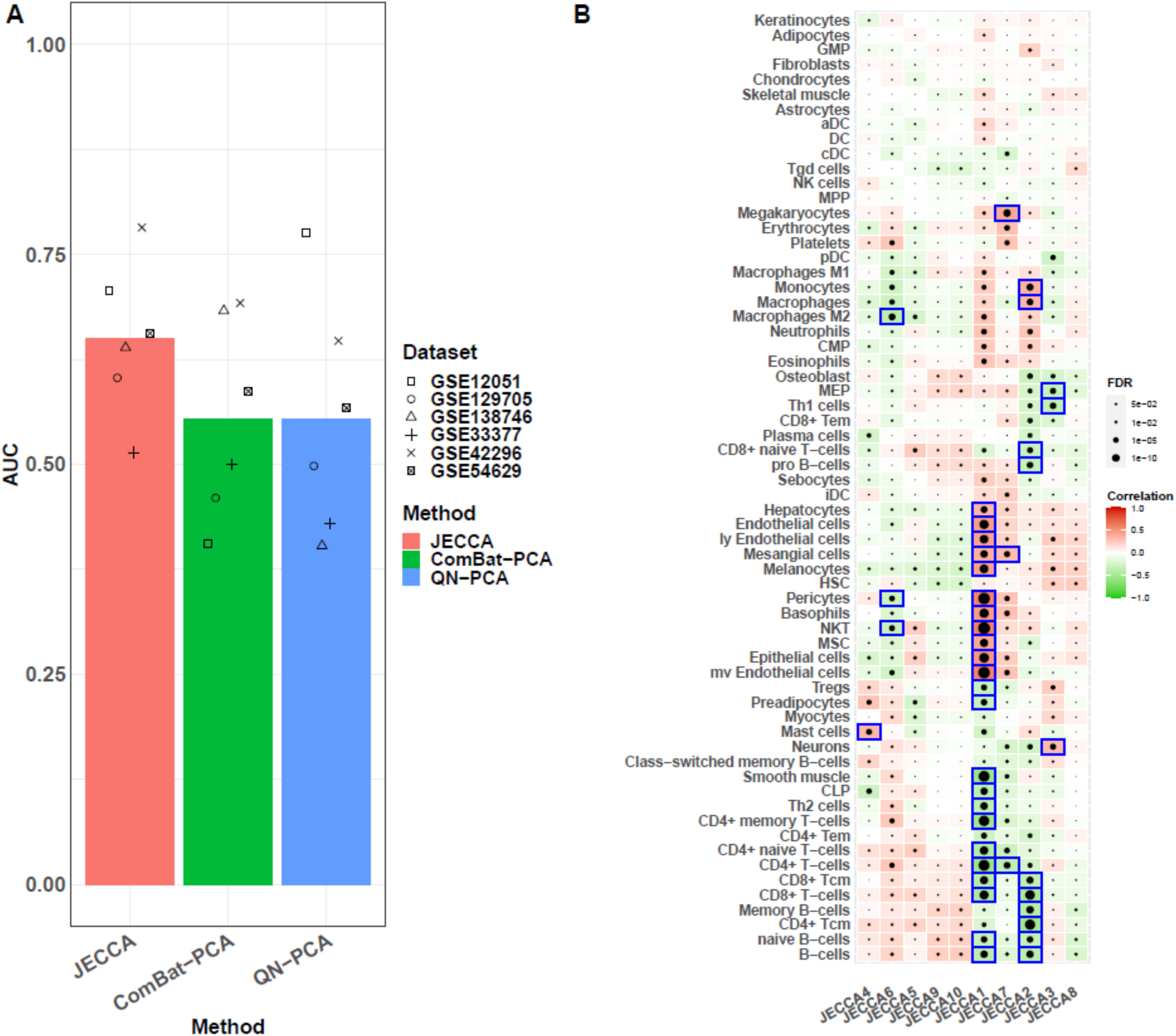
RA classification and correlation analyses. (A) Comparison of JECCA, ComBat-PCA, and QN-PCA methods by classification analysis on six RA datasets. In classification analysis, the five RA datasets were used as training data and another one dataset was used as testing data. (B) Spearman’s correlation between JECCA derived embeddings and xcell predicted cell type enrichment scores. The enrichment scores were calculated based on the gene expression profile obtained from each JECCA embeddings. Blue brackets mark the absolute correlation greater than 0.3.

To uncover the biological information within each JECCA embedding, we performed cell type enrichment analysis on the expression data^47^. The correlation between the predicted cell type enrichment scores and each embedding revealed that JECCA2, the most significant embedding associated with responder/non-responder status (higher in responder shown in Supplementary Table 3), exhibited a negative correlation with T-cells and B-cells, while showing a positive correlation with monocytes (Figure 5B). Based on the weights of the JECCA2 embedding, we concluded that patients responding to anti-TNF therapies exhibited decreased T-cells and B-cells, and increased monocytes. This observation aligns with a recently published paper^43^, which identified an association between innate and adaptive cell populations and the response to anti-TNF therapy. JECCA also revealed that baseline mast cell signature in blood is associated with probability of therapeutic response to anti-TNF treatment in the fourth embedding. Mast cells are tissue-resident cells of the innate immune system and their role in RA has been well studied and the varied signature identified here might represent MC progenitors circulate in blood stream ^48,49^. Overall, our findings highlight the importance of variations in cell composition between anti-TNF drug responders and non-responders in the context of RA, which could be used to guide the treatment assignments to RA patients.

### JECCA Enables Cross Species Alignment

In drug discovery and validation, mouse models are commonly utilized due to their ease of manipulation in a laboratory environment, which allows researchers to control variables, perform targeted genetic modifications, and conduct precise experiments. It is critical to appreciate that these models do not fully replicate the complexity of human disease, as there can be inherent biological and drug response differences between species. Here we showed that JECCA is not only able to extract shared information across multiple datasets, but also enables alignment of data from different species to identify common biological features.

In this case study, we employed our method to demonstrate its benefits in cross-species alignment using UC data as an example. We initially considered a dextran sodium sulfate (DSS)-induced mouse model, which mimics the superficial inflammation observed in UC ^50^. In this study a total of 26 DSS mouse samples were collected over a span of 14 days, with the first 7 days involving DSS treatment followed by 7 days of recovery. The paper revealed conserved inflammatory pathways between mice and humans and successfully identified homologous genes in mice that facilitated clustering of human IBD patient samples into two groups highly associated with response and non-response status to IFX treatment. This finding supports the existence of shared biological information, particularly regarding the inflammatory process, between the DSS mouse model and human UC patients.

We applied JECCA to combine four human UC datasets with the DSS mouse dataset directly, aiming to identify embeddings that capture shared information. Subsequently, we conducted classification analyses using embeddings from the four human IBD datasets and the DSS mouse dataset as training and testing data, respectively. Interestingly, we observed a U-shaped probability of predicted responder in the mouse data, which decreased from day 0 to day 7 and then increased after day 7, mirroring the longitudinal experimental design of the DSS mouse model (Figure 6A). This U-shape pattern suggested that non-response in human data might exhibit more severe inflammation compared to responses. Moreover, when we performed the classification analysis in reverse, using human data as testing and DSS mouse data as training to transfer the mouse “day” labeling to human samples, we observed a high proportion of non-responders during days 4 to 10, further indicating more severe inflammation in non-responders (Figure 6B).

**Figure 6.**
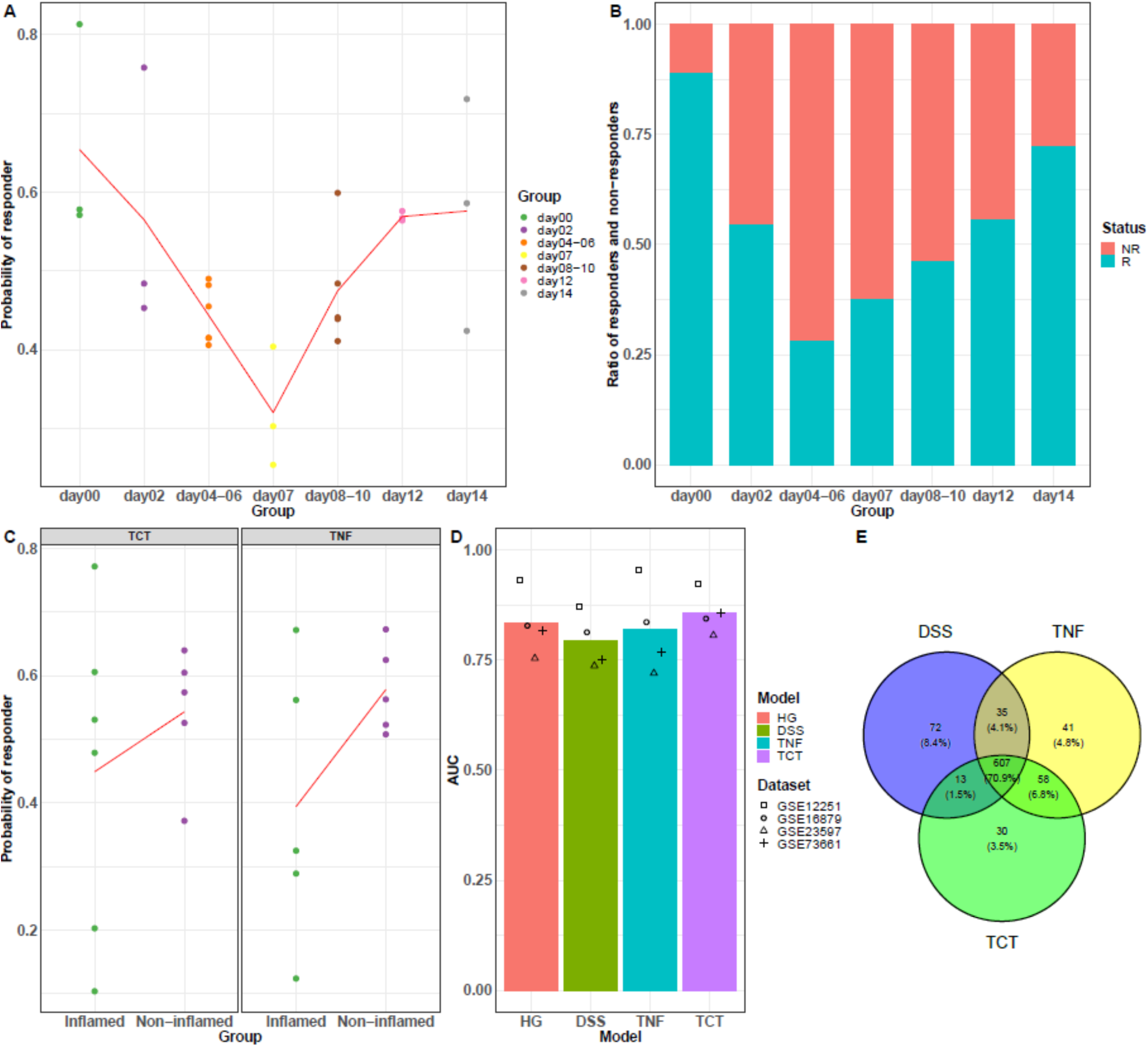
Mouse-human cross species alignment study. (A) Predicted probability of response to IFX for DSS mice. DSS mice are grouped by day information in the longitudinal experiment. Red line represents the average probability in each group. The predicted probability are estimated in a classification analysis using four IBD human datasets as training data and DSS mouse dataset as testing data (B) Barplots of proportion of human responders and non-responders predicted by day information in the DSS mouse longitudinal experiment. The predicted day information are obtained in a classification analysis using DSS mouse dataset as training data and four IBD human datasets as testing data. (C) Predicted probability of response to IFX for TCT and TNFΔARE mice. The predicted probability are estimated in a classification analysis using the four IBD human datasets as training data and TCT or TNFΔARE mouse dataset as testing data. Red lines represent the average predicted probability of inflamed and non-inflamed mice. (D) Comparison of three mouse models and one human model by classification analyses on derived JECCA embeddings. In three mouse models, the JECCA embeddings were derived from the four human datasets combined with each of the three mouse model datasets. In classification analysis, only JECCA embeddings of the four IBD human datasets were used with three datasets as training data and another one dataset as testing data. (E) Venn Diagram of the number of significant reactomPA pathways in three mouse models.

The ability of JECCA to perform cross-species alignment allows for the comparison of different animal models, aiding in the selection of the most suitable models and timepoints for a specific research objective. Next, we considered datasets from three mouse models of intestinal/colon inflammation: the DSS-induced model, the TNFΔARE model^51^, and the T cell transfer (TCT) model^51^. Each model has its own advantages and limitations concerning IBD research. For example, the acute DSS model recapitulates early aspects of human ulcerative colitis (UC) but fails to represent the chronic nature of this disease^52^. Conversely, the TNFΔARE (TNF over expression) and TCT models represent chronic inflammation, more reminiscent of Crohn’s Disease^53^. We employed JECCA using the four human IBD datasets in combination with each of the three mouse model datasets, separately. By generating embeddings through JECCA, we performed a similar analysis to that described for the DSS mouse model. Initially, we observed a low probability of predicted responder in the inflamed mouse samples from both the TNFΔARE and TCT models, consistent with the findings from the DSS model (Figure 6C). Subsequently, we compared the three models alongside the four human IBD data model by performing a classification analysis on the embeddings created from the four human IBD datasets, with three datasets used as training data and the remaining dataset serving as the testing data for responder/non-responder prediction. The results (Figure 6D) revealed that the embeddings generated from the TNFΔARE and TCT models achieved better AUC metrics compared to the DSS model and the performance of including TCT model even surpassed that of the IBD 4 human data model. This might suggest that the chronic inflammation variations represented by TNFΔARE and TCT models contain more biological relevant information in line with human UC data to assess anti-TNF treatment. Further subsequent pathway analysis (Figure 6E) showed that JECCA can identify pathway signatures either shared between the models or unique to each of them. Figure 6E unveiled that all three mouse models shared >70% of the identified human IBD pathways. The TNFΔARE and TCT models shared more pathways than with the DSS model.

Among these shared pathways (Supplementary Table 3), many of them are known to be related to chronic inflammation disease progression such as, Bile acid and bile salt metabolism^54^, RIP-mediated NF-kB activation via ZBP1^55^ and ABC transporters in lipid homeostasis^56^. The ability to align data from different species using JECCA allows researchers to find shared and distinct molecular processes between human and different preclinical models, enabling exploration of conserved pathways, genes, and mechanisms that contribute to disease development and progression. This cross-species alignment helps bridge the gap between animal models and human diseases, enhancing the predictive power of preclinical studies and facilitating the translation of findings to human clinical applications.

## Discussion

Incorporation of multiple datasets into a single analysis allows researchers to capture a broader perspective of disease biology and mitigates the limitations of individual datasets. Integration of diverse -omics datasets, such as animal model data, cell-line compound screening data, and human patient data, enables researchers to gain insights into biological mechanisms which are impossible to do in human alone, which is essential in genomic medicine development. However, such data integration remains a key challenge in systems biology and drug development.

More importantly, the application of JECCA and similar analytical approaches enables researchers to make more informed decisions in selecting appropriate animal models that closely mimic key biological pathways present in human diseases, thereby improving the translational relevance of preclinical studies. Animal models are critical components of drug development as they are needed to evaluate pharmacology of potential drugs. By enhancing our understanding of disease models, researchers can accelerate the development of effective therapies and improve patient outcomes.

To address the heterogeneity of the datasets, including the differences in platforms or species, we first identified the consensus genes shared across all datasets for JECCA and downstream pathway analysis. During the JECCA application, we applied row and column scaling to mitigate the influence of technique effects, species-specific variations, and outliers. Incorporating this scaling strategy in JECCA showed a clear advantage over the quantile normalization method, which was proposed in a review study^46^ to handle datasets with the mixture of microarray and RNA-seq data.

We have presented JECCA, a novel method to generate embeddings that capture shared biological variations across datasets. JECCA offers a powerful tool for data integration and comparative analysis. We have demonstrated that JECCA embeddings can be used to perform clustering, classification, and prediction analyses, to unravel important disease-related features and identify potential biomarkers or therapeutic targets. They can also be used to build translational models to align human and mouse data. We envision other applications of JECCA, such as cross disease gene expression data integration looking for common biological features, cell line and tissue data integration, predictive biomarker identification by modeling placebo and treatment arms together.

While JECCA has demonstrated broad applicability across various biological studies, there are some limitations worth noting. First, JECCA can be time-consuming with large datasets. For instance, in our breast cancer study involving around 1,000 subjects, the analysis took approximately 20 minutes to complete. Therefore, for multiple datasets with a large number of subjects, JECCA may require several hours to finish the analysis. Second, JECCA cannot fully eliminate technical effects or uninteresting biological variables (e.g. age), which means some of the generated embeddings may be associated with these factors. If the impact of such uninteresting variables dominates the datasets, JECCA may struggle to identify embeddings associated with the variables of interest.

JECCA is a statistical method designed to identify linear associations across multiple datasets. The performance of JECCA may be further improved through incorporating nonlinear CCA^57^ or Deep Variational CCA^58^, which is an important direction for future work. Aditionally, with the advancements in high-throughput proteomics techniques like Luminex, Simoa, and Olink, there is a promising opportunity to extend JECCA to proteomics data, which more closely reflects biological reality compared to transcriptomics.

## Methods

### PCA, CCA and JECCA

#### 1. Conventional PCA and CCA

Principle Components Analysis (PCA) is a method that finds linear combinations within a data set with the goal of maximizing the amount of variation that is explained by those Principal Components.

In PCA, given a standardized dataset *X* with n sample and *p* variables, the principal component 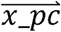 (projected values) is given by the linear combination of the original variables 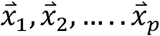 (default is column vector, scaled with zero mean and unit variance), *where i* = 1, … .*p*., and 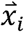 is a *n* * 1 column gene vector indicating the expression vector of gene 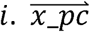 is a linear combination of gene vectors, which can be written as 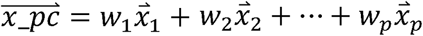.

In order to maximize variance, the first weight vector 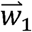 has to satisfy:

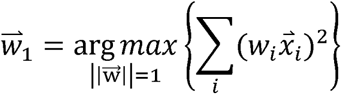

The first principal component PC1 represents the component that retains the maximum variance of the data. The *2nd* component will then be the leading component of the residuals after subtracting the first principal component from matrix *X* and so on. Principal components may be considered metagenes, aggregate pattern of gene expression serving as surrogates of specific biomolecular events or pathways which are likely to be found associated with a phenotype.

Where PCA focuses on finding linear combinations that account for the most variance in one data set, Canonical Correlation Analysis focuses on finding linear combinations that account for the maximal correlation in two datasets. CCA has been extensively used in multi-omics datasets integration to extract latent features shared between multiple assays, e.g., transcriptome, proteomics, and genomics. In this case, we have multiple assay measurements on the same set of subjects, the so-called vertical integration.

#### 2. Joint-Embedding based on Canonical Correlation Analysis (JECCA)

JECCA is designed to integrate independent gene expression data across studies on the same set of variables (genes), the so-called horizontal integration. We describe the two datasets integration scenario first which can be easily extended to multi datasets analysis. Now let’s consider two datasets, *X* with *n* sample and *p* variables and *Y* with m sample and *p* variables, e.g., the same set of genes measured on two different sets of subjects. 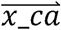 is a linear combination of sample vectors in *X*, which can be written as 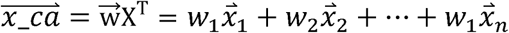, and contrary to the classical PCA, here 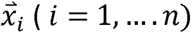 is a *p*\*1 column sample vector and 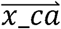 is called a meta-sample. For *Y* dataset (*p* genes with *m* samples), we have 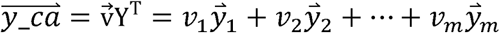, *where* 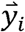, with *i* = 1, …, *m,* is also a *p*\*1 sample vector.

JECCA seeks vectors 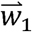 and 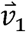 such that the random variables 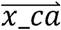 and 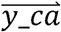 maximize the correlation:

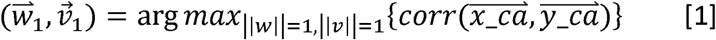

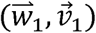 is the first pair of canonical variables. In computing the *r*th canonical pair, the orthogonality among the canonical variate pairs must hold^23^. The elements in 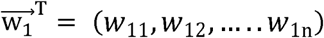 and 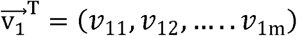 are the sample level embeddings for dataset *X* and *Y*, respectively, representing the shared signal between *X* and *Y*. The estimated aligned embeddings (Figure 1) facilitate many downstream analyses, e.g., disease endotype identification. Samples which are similar at the molecular level should demonstrate similar or more correlated embeddings. We adopted the penalized matrix decomposition approach implemented in R package, PMA, for computing the CCA embeddings^59,60^. For integrating more than two data sets, the objective function in [1] is extended to maximize the sum of all pairwise correlations of the canonical variables^60^.

### Datasets preprocessing

The datasets used in four case studies are summarized in Supplementary Table 1. For the breast cancer study, we collected 11 GEO datasets with a total of 1604 samples. The preprocessed data was downloaded from a recently published paper^24^. We performed log2 transformation on the data before downstream analysis. In the UC study, we collected 5 GEO datasets with a total of 345 baseline samples. The same preprocessing steps were conducted for these 5 datasets as described in a relevant paper^50^. In the RA blood study, we collected 6 GEO datasets with a total of 275 baseline samples. For microarray data, we obtained the preprocessed log2 transformed data using the “getGEO” function in the GEOquery Bioconductor package. As for RNA-seq data, we downloaded the raw counts data from the NCBI GEO database. The counts were normalized using the TMM^61^ normalization method and further processed into log2 transformed counts per million (CPM) data for downstream analysis. In the cross-species alignment study, we downloaded two RNA-seq mouse datasets from the NCBI GEO database. In the DSS model, we obtained the processed log2 transformed CPM data from the NCBI GEO database. For the TNFΔARE and TCT models, we downloaded raw counts data from the NCBI GEO database and applied the same RNA-seq preprocessing steps as described in the RA study to obtain log2 transformed CPM data for downstream analysis.

### JECCA, ComBat-PCA, and QN-PCA analysis

Firstly, we conducted gene-level scaling on each individual dataset. Next, we merged the multiple datasets within each case study by selecting the common genes present in all the datasets. For the cross-species alignment, only orthologues genes that appeared in both human and mouse datasets were utilized. We applied JECCA to merged data to extract embeddings. In the ComBat-PCA method, we treated each GEO dataset as a batch and performed batch correction using the ‘ComBat’ function from the SVA Bioconductor package. On the other hand, in the QN-PCA method, we employed the ‘normalize.quantiles.use.target’ method from the preprocessCore Bioconductor package. For normalization, we utilized the log2-transformed microarray data from the GEO33377 dataset as the reference data. Subsequently, we performed PCA analysis on the batch-corrected data from the ComBat method or the QN-normalized data. This analysis allowed us to obtain principal components, which served as embeddings for downstream clustering or classification analysis.

### Unsupervised clustering, differential gene expression and classification analysis

To conduct unsupervised clustering on the JECCA-derived embeddings, we utilized the ‘FindClusters’ function from the Seurat Bioconductor package. The default Louvain clustering method was employed. For dimension reduction and visualization of the embeddings, we applied the ‘RunUMAP’ function from Seurat.

For the human UC study, differential gene expression analysis was performed between the two identified clusters using ‘eBayes’ function in limma Bioconductor package. Genes with fold change greater than 2 and FDR less than 0.05 were selected as differential expressed genes. The downstream pathway analysis of up- and down-regulated genes were performed by Reactome enrichment analysis using ‘enrichPathway’ function from clusterProfiler bioconductor package.

For the classification analysis, we utilized the ‘ranger’ function from the R package ranger to perform random forest on the training data. Subsequently, we used the ‘predict’ function to make predictions on the testing data. The training and testing data were based on the embeddings obtained from the JECCA, ComBat-PCA, or QN-PCA method.

In each disease study, we employed the ‘auc’ function from the precrec R package to calculate the Area Under the Curve (AUC) scores. For three-class classification evaluation in breast cancer grade application, we utilized the ‘VUS’ function from the DiagTest3Grp R package to calculate the Volume Under a three-class ROC Surface (VUS).

### GSEA pathway, ssGSEA and cell type enrichment analysis

For the JECCA method, we utilize embeddings and gene expression profiles of each dataset to generate meta-samples, a linear combination of sample vectors weighted by the embeddings. To identify the most significant pathways of the meta-samples per dataset, we conducted GSEA analysis. For each JECCA derived embedding, we obtain the list of significant pathways per dataset. Subsequently, we employ the ‘AWFisher_pvalue’ function from the AWFisher R package to calculate a meta-p-value of the pathways across multiple datasets, enabling us to further select the most significant pathways across all datasets. We leveraged two Bioconductor packages: ‘clusterProfiler’ and ‘fgsea’ to perform GSEA pathway enrichment analysis using the ‘enrichGO’ and ‘fgsea’ function, respectively. The pathway databases are based on ‘Cellular component’ Gene Ontology annotation terms and ReactomePA.

For human UC study, we calculated ssGSEA enrichment scores for each dataset using ‘gsva’ function form the GSVA bioconductor package. The pathway databases are based on ReactomePA.

In cell type enrichment analysis, the enrichment scores were estimated for each sample using the ‘xCellAnalysis’ function from the xCell Bioconductor package.

## Data availability

All the datasets analyzed in this paper are available at https://github.com/xchen004/JECCA/tree/main/Data. JECCA function is available at https://github.com/xchen004/JECCA/blob/main/JECCA.R. R scripts to reproduce figures are available at https://github.com/xchen004/JECCA/tree/main/script.

## Conflicts of Interest

This manuscript was sponsored by AbbVie. AbbVie contributed to the design, research, and interpretation of data, writing, reviewing, and approved the content. XC, KS and YB are employees of AbbVie Inc. All authors may own AbbVie Stock.

## Contributions

YB and XC conceived the method and strategy. XC developed the coded algorithm and performed data analysis. YB and KMS developed the immunological applications of the method. XC, KMS and YB wrote the manuscript.

## Supplementary Figures and Tables

**Supplementary figure 1.**
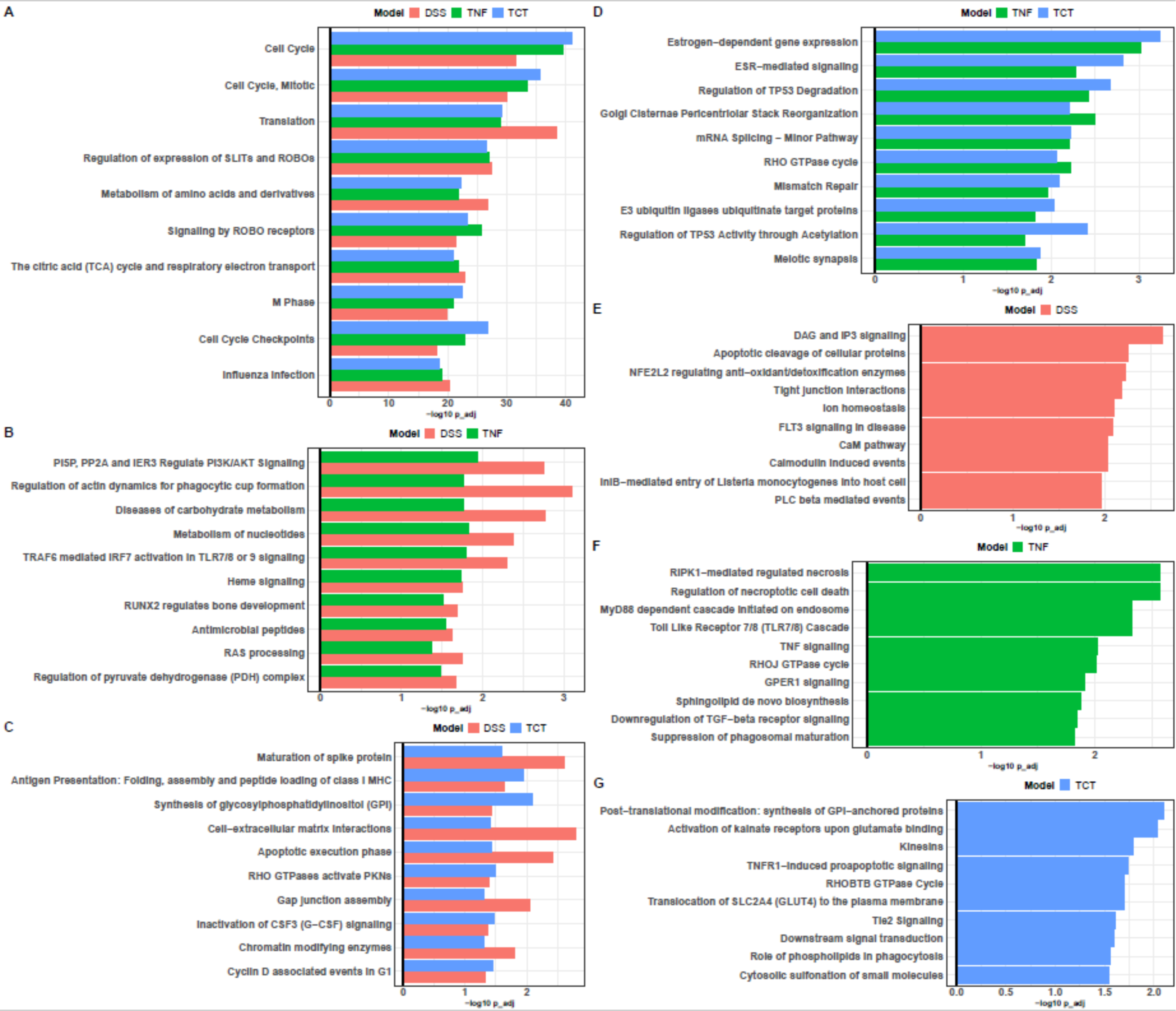
Top 10 significant reactomPA pathways in (A) all three mouse models, (B) DSS and TNF models, (C) DSS and TCT models, (D) TNF and TCT models, (E) DSS model, (F) TNF model, and (G) TCT model. (A-D) represents the overlapping pathways of two/three mouse models. (E-G) represents the exclusive pathways of each mouse model.

**Sup table 1:**
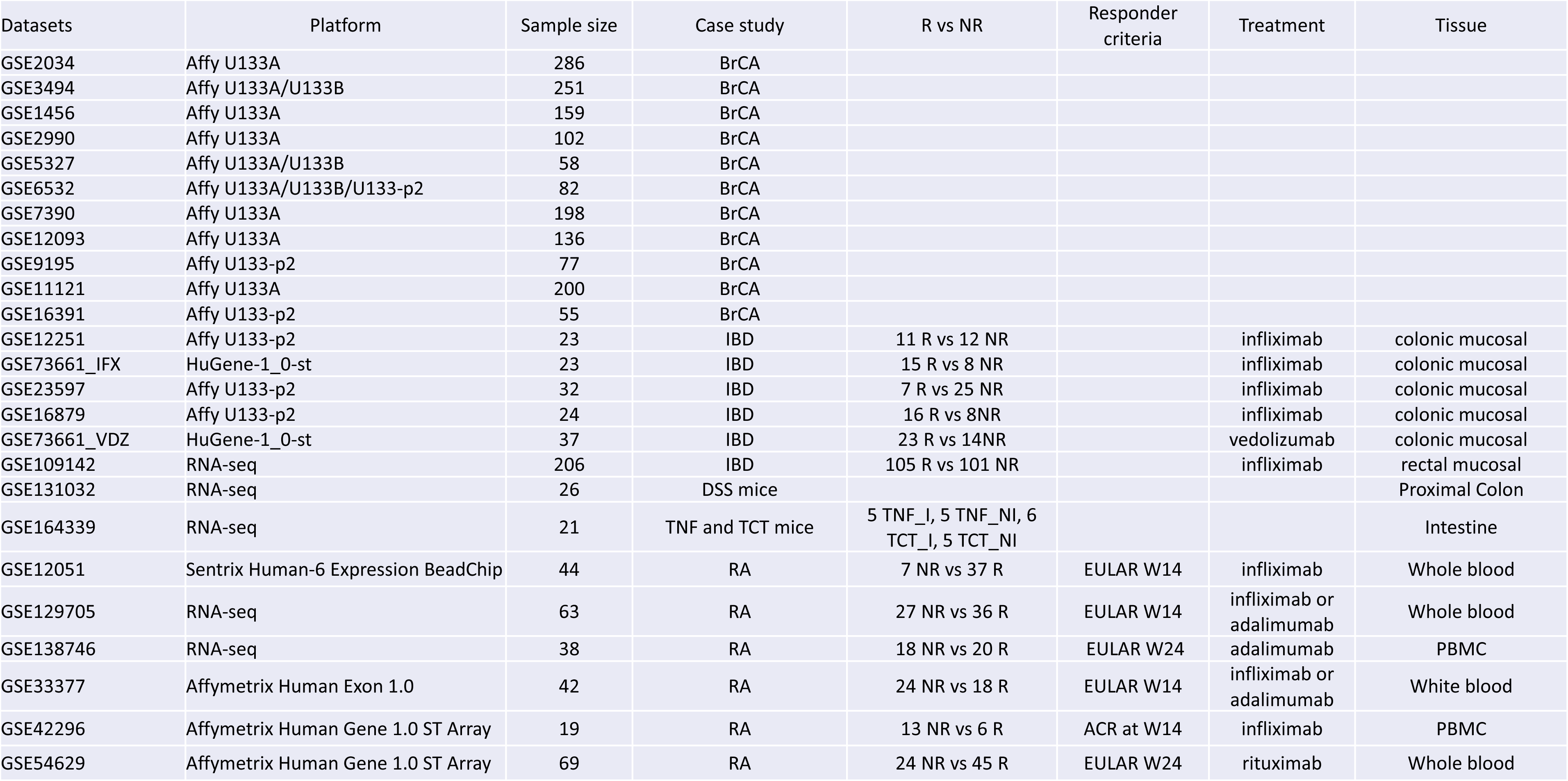

| Datasets | Platform | Sample size | Case study | R vs NR | Responder criteria | Treatment | Tissue |
| --- | --- | --- | --- | --- | --- | --- | --- |
| GSE2034 | Affy U133A | 286 | BrCA |  |  |  |  |
| GSE3494 | Affy U133A/U133B | 251 | BrCA |  |  |  |  |
| GSE1456 | Affy U133A | 159 | BrCA |  |  |  |  |
| GSE2990 | Affy U133A | 102 | BrCA |  |  |  |  |
| GSE5327 | Affy U133A/U133B | 58 | BrCA |  |  |  |  |
| GSE6532 | Affy U133A/U133B/U133-p2 | 82 | BrCA |  |  |  |  |
| GSE7390 | Affy U133A | 198 | BrCA |  |  |  |  |
| GSE12093 | Affy U133A | 136 | BrCA |  |  |  |  |
| GSE9195 | Affy U133-p2 | 77 | BrCA |  |  |  |  |
| GSE11121 | Affy U133A | 200 | BrCA |  |  |  |  |
| GSE16391 | Affy U133-p2 | 55 | BrCA |  |  |  |  |
| GSE12251 | Affy U133-p2 | 23 | IBD | 11 R vs 12 NR |  | infliximab | colonic mucosal |
| GSE73661_IFX | HuGene-1_0-st | 23 | IBD | 15 R vs 8 NR |  | infliximab | colonic mucosal |
| GSE23597 | Affy U133-p2 | 32 | IBD | 7 R vs 25 NR |  | infliximab | colonic mucosal |
| GSE16879 | Affy U133-p2 | 24 | IBD | 16 R vs 8NR |  | infliximab | colonic mucosal |
| GSE73661_VDZ | HuGene-1_0-st | 37 | IBD | 23 R vs 14NR |  | vedolizumab | colonic mucosal |
| GSE109142 | RNA-seq | 206 | IBD | 105 R vs 101 NR |  | infliximab | rectal mucosal |
| GSE131032 | RNA-seq | 26 | DSS mice |  |  |  | Proximal Colon |
| GSE164339 | RNA-seq | 21 | TNF and TCT mice | 5 TNF_I, 5 TNF_NI, 6 TCT_I, 5 TCT_NI |  |  | Intestine |
| GSE12051 | Sentrix Human-6 Expression BeadChip | 44 | RA | 7 NR vs 37 R | EULAR W14 | infliximab | Whole blood |
| GSE129705 | RNA-seq | 63 | RA | 27 NR vs 36 R | EULAR W14 | infliximab or adalimumab | Whole blood |
| GSE138746 | RNA-seq | 38 | RA | 18 NR vs 20 R | EULAR W24 | adalimumab | PBMC |
| GSE33377 | Affymetrix Human Exon 1.0 | 42 | RA | 24 NR vs 18 R | EULAR W14 | infliximab or adalimumab | White blood |
| GSE42296 | Affymetrix Human Gene 1.0 ST Array | 19 | RA | 13 NR vs 6 R | ACR at W14 | infliximab | PBMC |
| GSE54629 | Affymetrix Human Gene 1.0 ST Array | 69 | RA | 24 NR vs 45 R | EULAR W24 | rituximab | Whole blood |

**Sup table 2:**
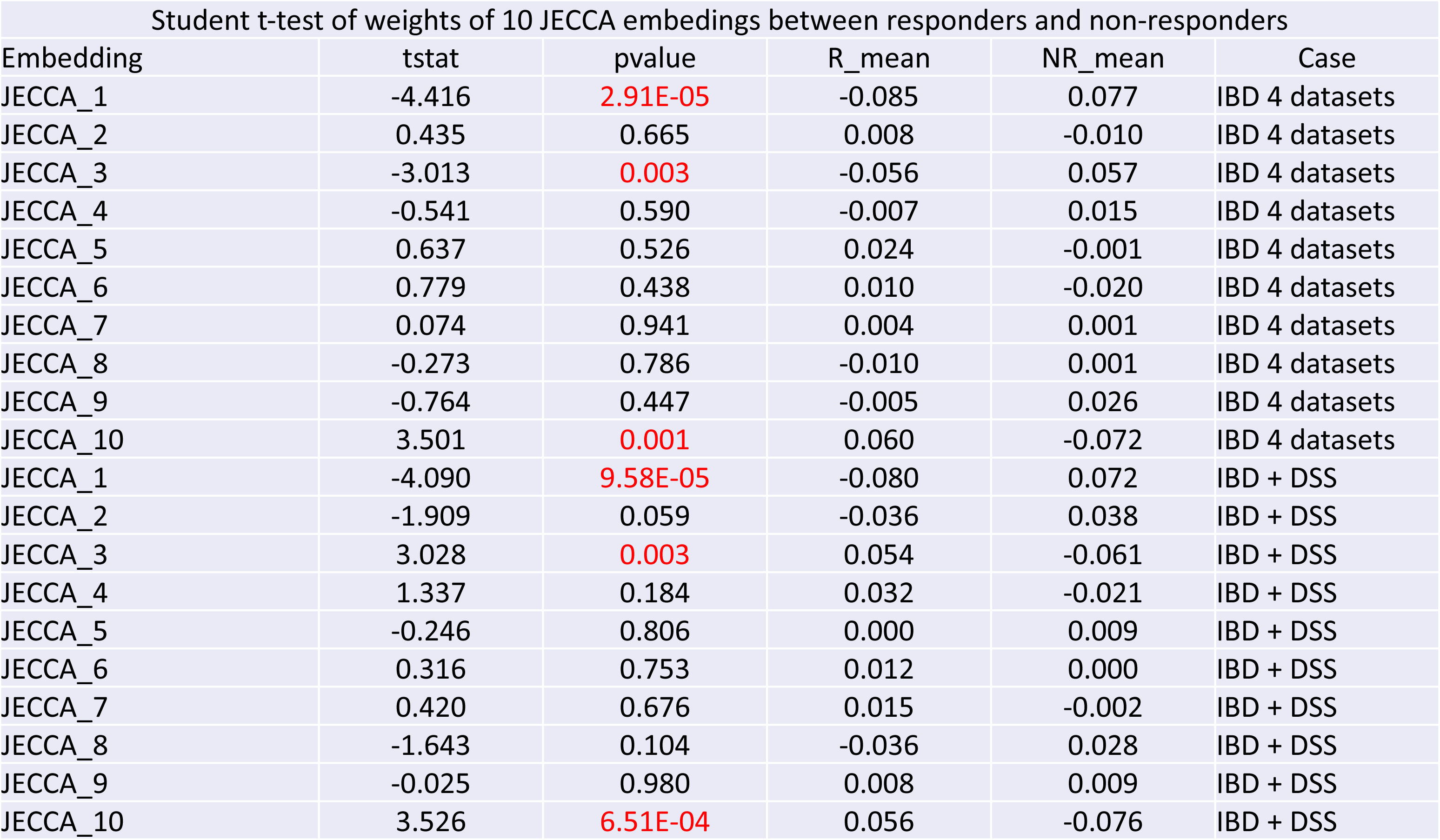
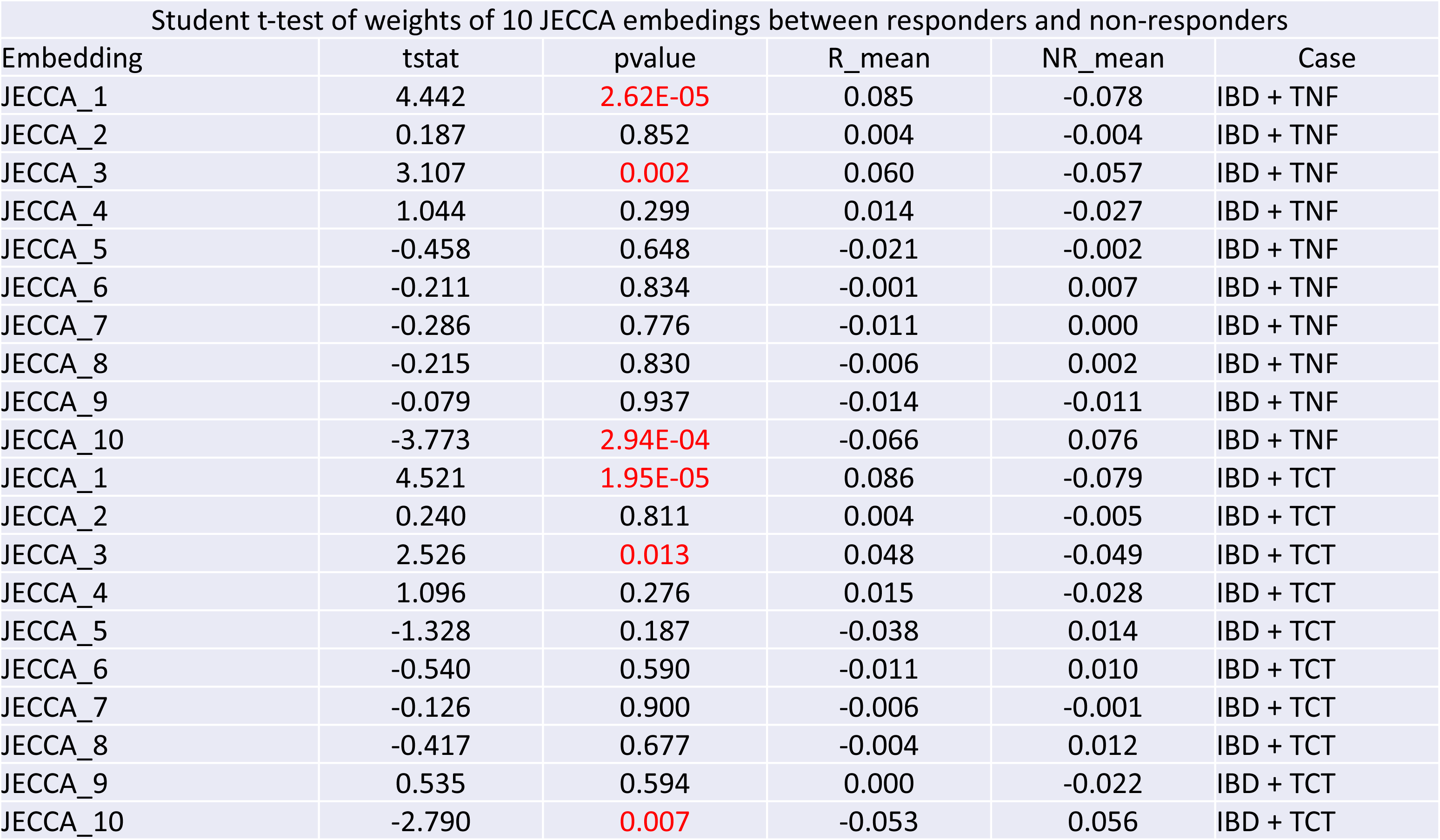
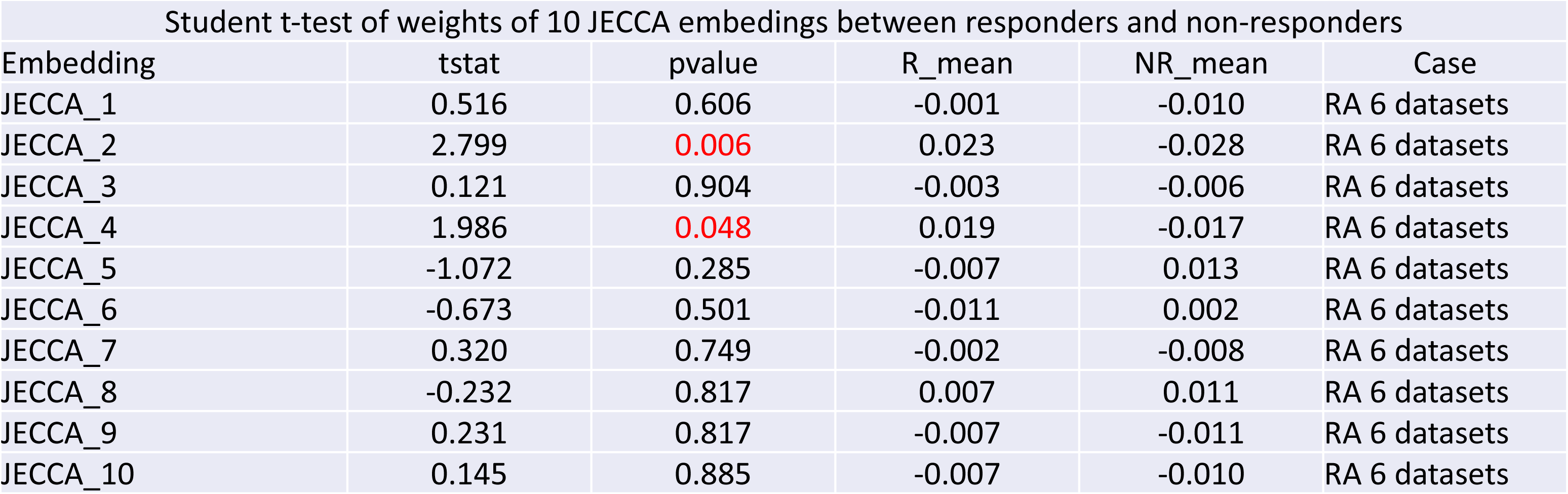

## Notes

https://github.com/xchen004/JECCA

### Competing Interest Statement

The authors have declared no competing interest.

